# A CD57^+^ cytotoxic CD8 T cell subset associated with fibrotic lung disease in systemic sclerosis

**DOI:** 10.1101/2025.01.27.635121

**Authors:** Takanori Sasaki, Ye Cao, Kathryne Marks, Kazuhiko Higashioka, Mehreen Elahee, Richard Ainsworth, Kim Taylor, Nunzio Bottini, Paul Wolters, Mari Kamiya, Edy Y. Kim, Francesco Boin, Deepak A. Rao

**Author notes:** Correspondence Francesco Boin, MD, Division of Rheumatology, Cedars-Sinai Medical Center, 8700 Beverly Blvd, B 122, Los Angeles, CA 90048,; Deepak Rao, MD PhD, Division of Rheumatology, Inflammation, Immunity, Brigham and Women’s Hospital, Hale Building for Transformative Medicine, 6002R, 60 Fenwood Road, Boston, MA, USA. Drs Boin and Rao contributed equally to this work.

## Abstract

Interstitial lung disease (ILD) is a major cause of morbidity and mortality in systemic sclerosis (SSc); however, the immunopathologic mechanisms driving lung disease in SSc are unclear. T cells have been implicated as a likely driver of lung injury in SSc. Here, we have evaluated T cells in blood and lungs of patients with SSc-ILD and identified a specific population of cytotoxic CD8 T cells that is expanded in SSc-ILD patients. Cytotoxic effector memory CD8 T cells marked by CD57 expression are preferentially expanded in SSc-ILD patients compared to SSc patients without ILD and controls and show prominent clonal expansion. These CD57^+^ T effector memory (TEM) cells differ from T effector memory cells re-expressing CD45RA (TEMRA) transcriptomically and functionally, with cytotoxic function that is enhanced by CD155 engagement of the costimulatory receptor CD226. Analyses of cells from ILD lungs indicate endothelial cells as a likely source of CD155 to activate CD57^+^ cytotoxic T cells. Together, the results implicate a CD57^+^ cytotoxic CD8 T cell population as a potential mediator of lung injury in SSc-ILD.

## Introduction

Systemic sclerosis (SSc) is a systemic autoimmune disease characterized by hardening of skin, fibrosis, and vascular endothelial damage (*1*). The disease can impact vital organs such as lungs, heart, and kidneys. Interstitial lung disease (ILD) is a frequent complication, occurring in approximately 40-75% of patients, and is associated with poor prognosis (*1*). Mycophenolate mofetil (*2*), cyclophosphamide (*3*, *4*), rituximab (*5*), tocilizumab (*6*), and nintedanib (*7*) have shown some benefit in SSc-ILD (*8–10*); however, the efficacy of these treatments remains limited, and SSc-ILD is still a leading cause of SSc-related mortality (*11*), underscoring the need for novel therapeutic approaches.

SSc-ILD pathology is characterized by irreversible and progressive pulmonary fibrosis. Early detection can improve prognosis; thus, current guidelines recommend for screening for ILD using pulmonary function tests (PFT), high-resolution CT (HRCT), and Scl-70 antibody testing, as well as monitoring ILD progression by PFT and HRCT after initiating SSc-ILD treatment (*8*– *10*). These metrics primarily evaluate pulmonary function decline and the extent or progression of lung fibrosis after it has already occurred, but do not quantify the magnitude of the ongoing immune-mediated injury. Delineation of such immunologic drivers in the peripheral blood of patients with SSc-ILD could aid the prompt identification of patients at risk for more severe and progressive SSc-ILD, allow measurement of the extent of the pathologic immune activation, and define with greater precision the efficacy of the ongoing treatments in suppressing potential immune-mediated injury.

Several studies have implicated T cells as drivers of tissue injury in SSc. Both CD4 and CD8 cytotoxic T lymphocytes (CTLs) accumulate within skin of patients with SSc and induce apoptotic death in dermal endothelial cells, potentially leading to tissue damage and fibrosis (*12*). A PD-1^hi^ cytotoxic CD4 T cell population is also expanded in the circulation of SSc-ILD patients (*13*). In addition, IL-13^+^ CD8 T cells are increased in the blood and skin of SSc patients, and their Th2-like functions contribute to skin fibrosis (*14*, *15*). Moreover, a recent single-cell RNA sequencing (scRNA-seq) study using lung samples from SSc-ILD has revealed an increase in CD8 tissue-resident memory T cells (*16*). However, the features of pathologic T cell activation that are most prominently activated in patients with SSc-ILD remain incompletely defined. In this study, we have generated extensive mass cytometry and scRNA-seq data of circulating lymphocytes from patients with SSc and sought to identify T cell phenotypes associated with the presence and severity of ILD. We found that CD57^+^ effector memory (CD57^+^ TEM) CD8 T cells are increased in patients with SSc-ILD and associated with ILD severity. This T cell subset shows substantial oligoclonality and accumulates within affected areas of fibrotic lungs. Their cytotoxic function is enhanced by activation of surface CD226, potentially by interaction with CD155 expressed on endothelial cells. These findings highlight a prominent cytotoxic pathway mediated by CD57^+^ TEM CD8 T cells associated with ILD in SSc patients and implicate the CD155-CD226 axis as a possible mediator of their activation and function.

## Results

### CD57^+^ TEM cells are expanded and associated with disease severity in SSc-ILD

To identify disease-associated changes in circulating lymphocytes in patients with SSc-ILD, we evaluated peripheral blood mononuclear cells (PBMC) from a cross-sectional cohort including 53 SSc-ILD patients, 29 SSc patients without ILD, and 18 healthy control patients without known autoimmune disease. In this cohort (Cohort 1), patients were mostly middle aged (54.5 ± 13.3 years) females (87%), predominantly White (60%), with variable skin involvement (diffuse cutaneous SSc in 39%). Detailed socio-demographic and clinical information are reported in Supplementary Table 1. We employed mass cytometry to analyze PBMC from these patients and focused our analyses on CD8 T cells given previous evidence supporting significant roles for this subset in SSc and ILD pathogenesis (*12*, *14–16*). To identify CD8 T cell phenotypes associated with SSc-ILD, we utilized covarying neighborhood analysis (CNA) (*17*), a cluster-free method to define disease-associated cell phenotypes in an unbiased manner. CNA analysis comparing CD8 T cells from SSc-ILD patients and controls identified a region of T cells significantly increased in SSc-ILD patients (FDR < 0.01, Figure 1A). This region contained cells marked by expression of CD57 and CD45RO and lacking CCR7, indicating a CD57^+^ effector memory (CD57^+^ TEM) phenotype. This population also exhibited high expression of T-bet and CX3CR1, two markers associated with CTLs, as well as activation markers CD38 and HLA-DR, and low expression of CD27 and CD56 (Supplementary Figure 1). Validation staining on a subset of samples by flow cytometry confirmed that CD57 and CD27 demonstrate an inverse expression pattern on CD8 TEM cells (Figure 1B).

**Figure 1.**
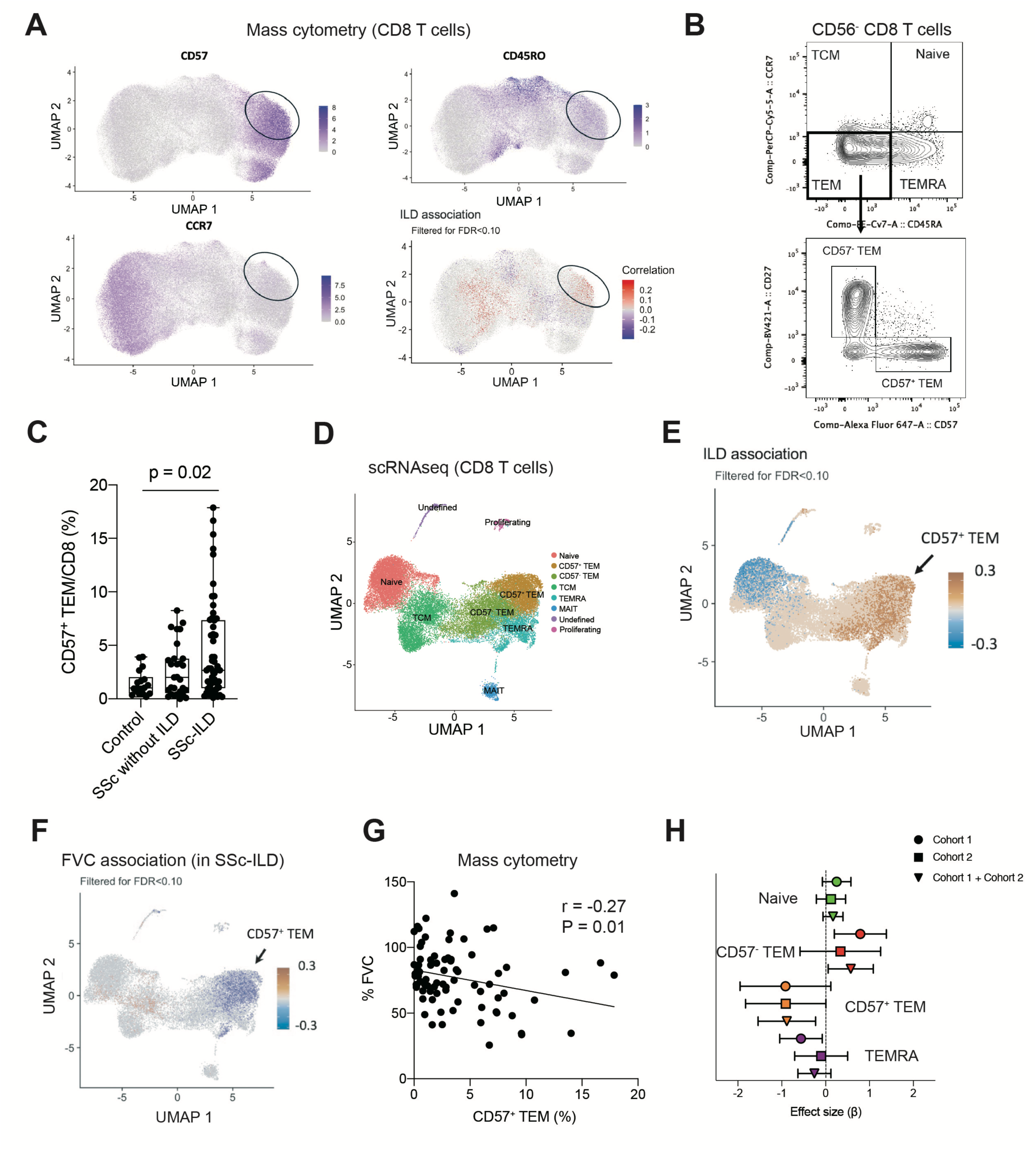
CD57^+^ TEM cells are expanded in the patients with SSc-ILD. **A.** CNA analysis of CD8 T cells from HC and SSc-ILD patients, adjusting for age and sex. Red indicates cell neighborhoods enriched in SSc-ILD patients. **B.** Flow cytometry detection of CD57 and CD27 in CD8 TEM cells. **C.** Quantification of CD57^+^ CD27^-^ CCR7^-^ CD45RO^+^ CD56^-^ cells among CD8 T cells (HC: n=18, SSc-non ILD: n = 29, SSc-ILD: n = 53). Kruskal-Wallis test with Dunn’s multiple comparison test. **D.** UMAP of CD8 T cell clusters in scRNA-seq dataset (SSc-non ILD: n = 29, SSc-ILD: n = 37) **E.** CNA analysis of CD8 T cells from SSc-non ILD and SSc-ILD, adjusting for age and sex. Red indicates cell neighborhood enriched in SSc-ILD patients. **F.** CNA analysis to assess correlation with FVC, adjusting for age and sex. **G.** Correlation analysis between CD57^+^ TEM cells and FVC. Spearman statistic shown. **H.** Effect size of CD8 T subset association with FVC by linear mixed-effect model. Mean ± 95% CI is shown. HC, healthy control.

Manual gating of CD8 T cells in the mass cytometry dataset confirmed that CD57^+^ CD27^-^ CCR7^-^ CD45RO^+^ CD8 T cells were significantly increased in SSc-ILD patients compared to controls but not in SSc patients without ILD (p = 0.02; control vs. SSc-ILD, mean: Control 1.5%, SSc without ILD 2.7%, SSc-ILD 4.5%, Figure 1C). To confirm expression of cytotoxic proteins in CD57^+^ TEM cells, we performed intracellular staining for granzymes using samples from 10 controls, 5 patients with SSc without ILD, and 5 patients with SSc-ILD. Flow cytometry analysis validated that almost 90% of CD57^+^ TEM express granzyme B. CD57^+^ TEM cells showed comparable levels of granzyme B to TEMRA, a well-known granzyme B^+^ CD8 T cell population (Supplementary Figure 2). The levels of granzyme B in CD57^+^ TEM cells were similar across controls, SSc-ILD, and SSc-without ILD groups, suggesting that high granzyme B expression is an intrinsic feature of CD57^+^ TEM cells.

To extend the characterization of expanded CD8 T cells in SSc-ILD patients, we analyzed PBMC from the same cohort of SSc patients by scRNA-seq and applied CNA analysis to the CD8^+^ T cells in the scRNA-seq data. CNA analysis demonstrated the enrichment of a similar cytotoxic CD8^+^ T cell population in SSc-ILD patients in the scRNA-seq dataset, highlighting a region of CD8 T cells with high expression of *GZMB* and *HNRNPLL*, a regulator of CD45RA splicing upregulated in CD45RO^+^ cells (Figure 1D, E, Supplementary Figure 3A). Although *B3GAT1* (encoding CD57) expression was not well detected in the RNA-seq data, the identified population lacked *CD27* expression, indicating that similar CTL population was identified in both mass cytometry and scRNA-seq analyses (Figure 1D, E, Supplementary Figure 3A).

To identify cell phenotypes associated with ILD severity, we used CNA to test for cell regions associated with forced vital capacity (FVC) among the SSc-ILD patients. This analysis again highlighted a region containing likely CD57^+^ TEM cells as associated with SSc-ILD severity, with a negative correlation between cells in this region and FVC (Figure 1F). In contrast, CNA analysis showed no association between CD57^+^ TEM cells and the modified Rodnan skin score (Supplementary Figure 4, FDR < 0.1), suggesting a specific association between CD57^+^ TEM cells and ILD. Manual gating of mass cytometry data confirmed a significant negative association between CD57^+^ TEM cells and FVC (Figure 1G). To further validate the association between CD57^+^ TEM cells and SSc-ILD severity, we quantified CD57^+^ TEM cells in samples in an independent cohort of 42 patients with SSc-ILD (Cohort 2) with similar clinical features to the initial cohort (Supplementary Table 1) using flow cytometry. A linear mixed-effects model assessing cytometry data from Cohort 1 and Cohort 2 confirmed a consistent negative association between CD57^+^ TEM cells and FVC across both cohorts (Figure 1H, Cohort 1 + Cohort 2, p = 0.009).

### CD57^+^ TEM cells are terminal effector cytotoxic CD8 T cells distinct from TEMRA

The above results indicated that CD57^+^ TEM cells are uniquely associated with SSc-ILD prevalence and severity in a pattern not shared with TEMRA cells. To further assess the transcriptomic signatures of CD57^+^ TEM cells and their differences from TEMRA cells, we performed bulk RNA sequencing of naive, CD57^-^ TEM, CD57^+^ TEM, and TEMRA CD8 T cells isolated from 5 SSc-ILD patients (Figure 2A). PCA analysis revealed that CD57^+^ TEM and TEMRA CD8 T cells are markedly distinct from naive and CD57^-^ TEM cells (Figure 2B). Naive CD8 T cells exhibited high expression of *CCR7*, *CD27*, *SELL*, *TCF7*, and *LEF1*. CD57^-^ TEM cells showed high expression of the early activation marker *CD69* and retained stem memory markers *TCF7* and *LEF1*, alongside evidence of proliferation indicated by MKI67 expression. CD57^-^ TEM cells also displayed high expression of *GZMK*, aligning with findings from flow cytometry and scRNA-seq analyses (Supplementary Figure 2, 3A). CD57^+^ TEM cells exhibited elevated expression of cytotoxic molecules and associated features, including *GZMA, GZMB*, *PRF1*, *ZEB2*, and *CX3CR1*, along with *TGFB1* (Figure 2C and Supplementary Figure 5). CD57^+^ TEM cells also showed high expression of *HLADRA* and *TNFRSF9* (encoding 4-1BB). TEMRA CD8 T cells also expressed high levels of cytotoxic molecules but uniquely demonstrated elevated expression of *IRF8*, *LAG3*, *TIGIT*, and *EOMES*. Other potentially pro-fibrotic cytokines, including *IL4*, *IL13*, and *TGFB2*, were expressed at low levels across all four populations (Supplementary Figure 5).

**Figure 2.**
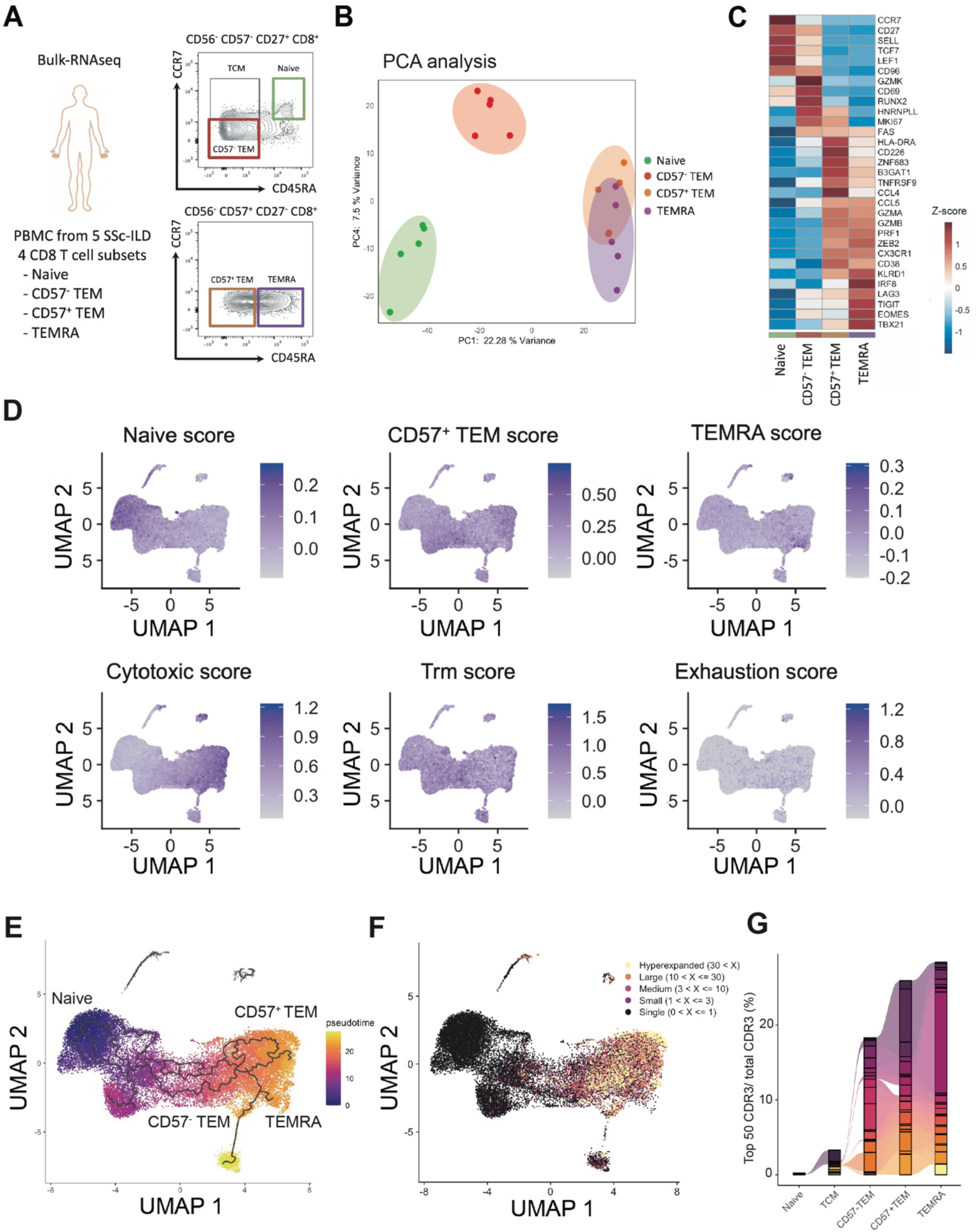
Developmental and clonal relationship between CD57^+^ TEM cells and TEMRA CD8 T cells. **A.** Sorting strategy of bulk RNA-seq for CD8 T cell populations in this study: naive CD8 T cells (CD57^-^ CD27^+^ CCR7^+^ CD45RA^+^ CD56^-^), CD57^-^ TEM cells (CD57^-^ CD27^+^ CCR7^-^ CD45RA^-^ CD56^-^), CD57^+^ TEM cells (CD57^+^ CD27^-^ CCR7^-^ CD45RA^-^ CD56^-^), and TEMRA CD8 T cells (CD57^+^ CD27^-^ CCR7^-^ CD45RA^+^ CD56^-^) were sorted from 5 SSc-ILD donors. **B.** PCA plots of naive CD8, CD57^-^ TEM cells, CD57^+^ TEM cells, and TEMRA CD8 T cells. **C.** Heatmap of expression of selected genes in bulk RNA-seq data from T cell subsets. **D.** Gene module scores in CD8 T cells in scRNA-seq data. **E.** Pseudotime analysis of CD8 T cells in the scRNA-seq dataset. **F.** Expanded TCR clones depicted onto the transcriptionally defined UMAP visualization. **G.** Proportions of top 50 clonotypes out of total clonotypes in naive, TCM, CD57^-^ TEM, CD57^+^ TEM, and TEMRA CD8 T cells.

Next, we interrogated expression of these bulk RNA-seq-derived signatures in the scRNA-seq profiling data from SSc patients to extend characterization of the cell populations associated with ILD. We established gene module scores for CD57^+^ TEM cells (44 upregulated genes) and for TEMRA cells (71 upregulated genes) (FDR < 0.1) from the bulk RNA-seq data (Figure 2D). In the scRNA-seq data, CD57^+^ TEM and TEMRA clusters demonstrated high levels of the respective gene module scores, confirming the distinct transcriptomic signatures of these cells across 2 transcriptional datasets. Cytotoxic gene signatures were highly enriched in the CD57^+^ TEM and TEMRA clusters, with no significant enrichment observed for tissue-resident memory CD8 T cells (Trm) or exhausted T cells. To investigate the developmental relationship between these subsets, we performed pseudotime analysis using Monocle 3 (*18*, *19*). The analysis revealed a differentiation trajectory beginning with naive CD8 T cells, transitioning through TCM and CD57^-^ TEM cells, and branching into both CD57^+^ TEM and TEMRA CD8 T cells (Figure 2E). T cell receptor (TCR) clonality analyses using TCR data captured in the scRNA-seq analysis demonstrated that both CD57^+^ TEM cells and TEMRA are highly clonally expanded in SSc patients (Figure 2F), and clonal overlap analysis showed that CD57^-^ TEM, CD57^+^ TEM, TEMRA, and proliferating clusters shared TCR repertoires (Supplementary Figure 3B), with identical TCRs by CDR3 sequence shared among CD57^-^ TEM, CD57^+^ TEM, and TEMRA CD8 T cells (Figure 2G). These findings indicates that cells within CD57+ TEM and TEMRA subsets can recognize a common antigen and share a developmental path, either through plasticity across the phenotypes or through a common progenitor state as CD57^-^ cells.

### CD226 augments the cytotoxic function of CD57^+^ TEM cells

To identify potential functional differences between CD57^+^ TEM and TEMRA CD8 T cells, we interrogated DEG between these two cell clusters in the scRNA-seq data (1,048 DEGs with FDR < 0.1, Figure 3A) and focused on genes encoding surface proteins and known regulators of T cell function. CD57^+^ TEM exhibited high expression of *ITGA4*, *ITGB1*, *CD226*, and *ZNF683* (encoding HOBIT), while TEMRA CD8 T cells showed elevated expression of *MX1*, *IFI44*, *IRF8*, and *TIGIT* (Figure 3A). Bulk RNA-seq and flow cytometry analyses confirmed that ITGA4 and ITGB1 (the two subunits of VLA4 integrin) were highly expressed in CD57^+^ TEM compared to TEMRA CD8 T cells (Figures 3B, 3C). In addition, CD226 and TIGIT were showed distinct patterns of expression between CD57^+^ TEM and TEMRA cells (Figure 3D, Figure 3E). Notably, the co-stimulatory receptor CD226 and the inhibitory receptor TIGIT share the same ligands, CD155 and CD112. Flow cytometry analysis verified that CD226 expression was higher in CD57^+^ TEM cells, while TIGIT expression was higher in TEMRA CD8 T cells, in both SSc-ILD patients and controls (Figures 3F, 3G). CD57^+^ TEM cells from the patients with SSc-ILD in Cohort 2 also showed a higher expression of CD226 and a lower expression of TIGIT compared to TEMRA cells (Figures 3H).

**Figure 3.**
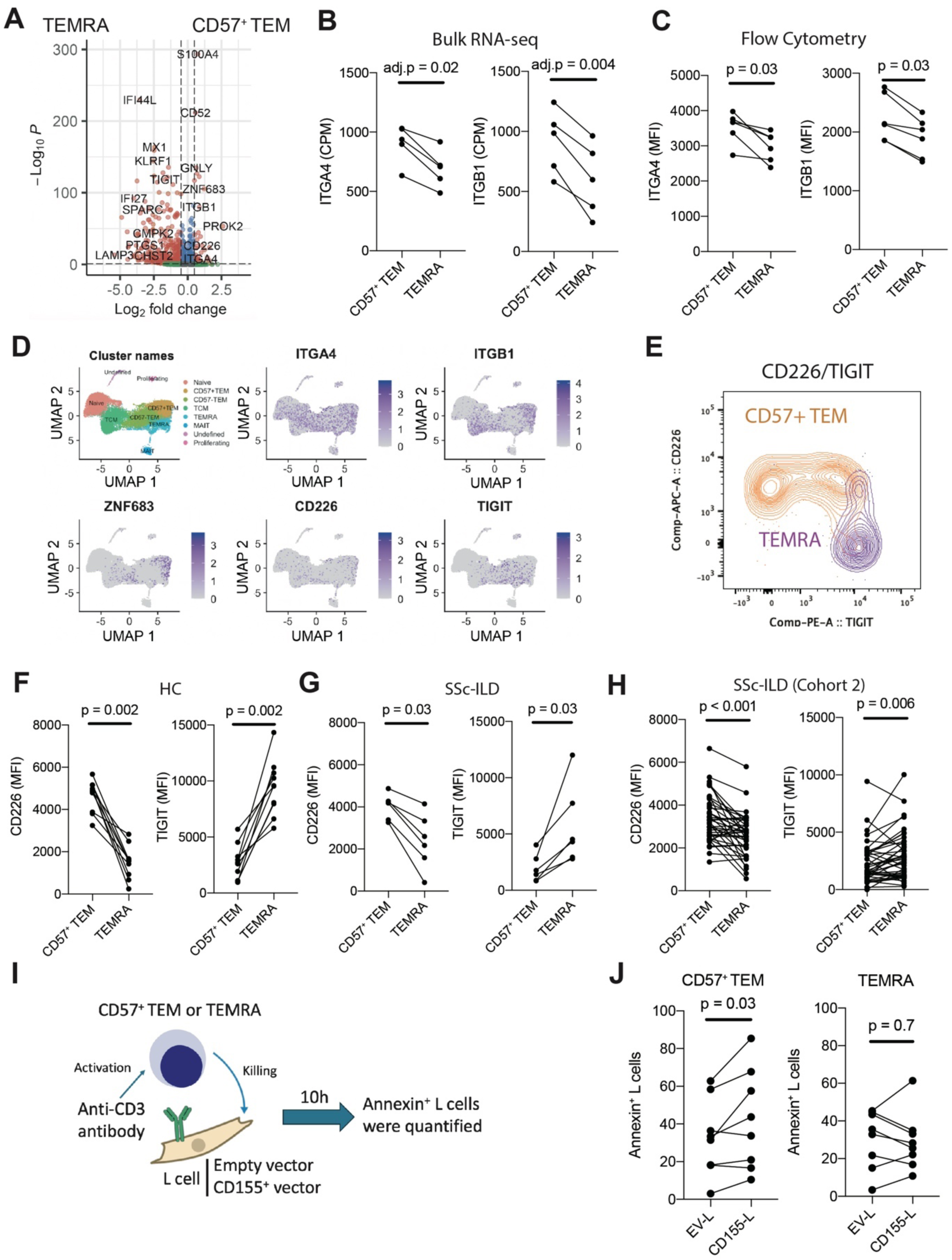
CD226 enhances the activation of CD57^+^ TEM cells. **A.** Differentially expressed gene analysis between the CD57^+^ TEM cluster and the TEMRA CD8 T cell cluster in scRNA-seq. **B.** Expression of *ITGA4* and *ITGB1* in CD57^+^ TEM and TEMRA CD8 T cells from patients with SSc-ILD assessed by bulk RNA-seq (n = 5) and flow cytometry (n = 6). **D.** Expression of *ITGA4*, *ITGB1*, *ZNF683*, *CD226*, and *TIGIT* in CD8 T cell clusters. **E.** Representative expression of CD226 and TIGIT on CD57^+^ TEM cells and TEMRA CD8 T cells by flow cytometry. **F, G.** CD226 and TIGIT expression on CD57^+^ TEM cells and TEMRA CD8 T cells by flow cytometry. The samples from 11 HC (F) and 6 SSc-ILD (G) were used. **H.** Validation of CD226 and TIGIT expression using Cohort 2 (SSc-ILD, n = 42). **I.** Protocol scheme of cytotoxicity assay to assess the effect of CD155 on CD57^+^ TEM and TEMRA CD8 T cells. **J.** Proportion of annexin^+^ L cells with or without CD155 after co-culture with CD8 T cell subsets. CD57^+^ TEM cells and TEMRA CD8 T cells were collected from 8 HC donors. Wilcoxon test shown. HC, healthy controls.

CD226 and TIGIT share the same ligands, CD155 and CD112, but can transmit opposing signals: CD226 provides activating signal, while TIGIT sends an inhibitory signal (*20*, *21*). Based on this, we hypothesized that the presence of CD155/CD112 may preferentially activate CD57^+^ TEM cells over TEMRA CD8 T cells. To evaluate the potential functional effects of CD155 on CD57^+^ TEM and TEMRA CD8 T cells, we applied a previously established cytotoxicity assay (*22*). CD57^+^ TEM or TEMRA CD8 T cells were co-cultured with murine fibroblast ‘L’ cells loaded with an anti-CD3 antibody to activate CD8 T cells and measure their killing activity (Figure 4I). To assess the specific effect of CD155, we compared the cytotoxicity induced by target L cells transduced to express human CD155 versus control transduced target L cells. CD57^+^ TEM demonstrated significantly increased cytotoxic activity when co-cultured with CD155^+^ L cells, whereas the cytotoxicity of TEMRA CD8 T cells was not altered (Figure 4J). These results suggest that CD57^+^ TEM cells differ from TEMRA in their regulation by CD155, with CD155 providing a stimulating effect on the cytotoxic function on the CD57^+^ TEM population expanded in SSc-ILD patients.

**Figure 4.**
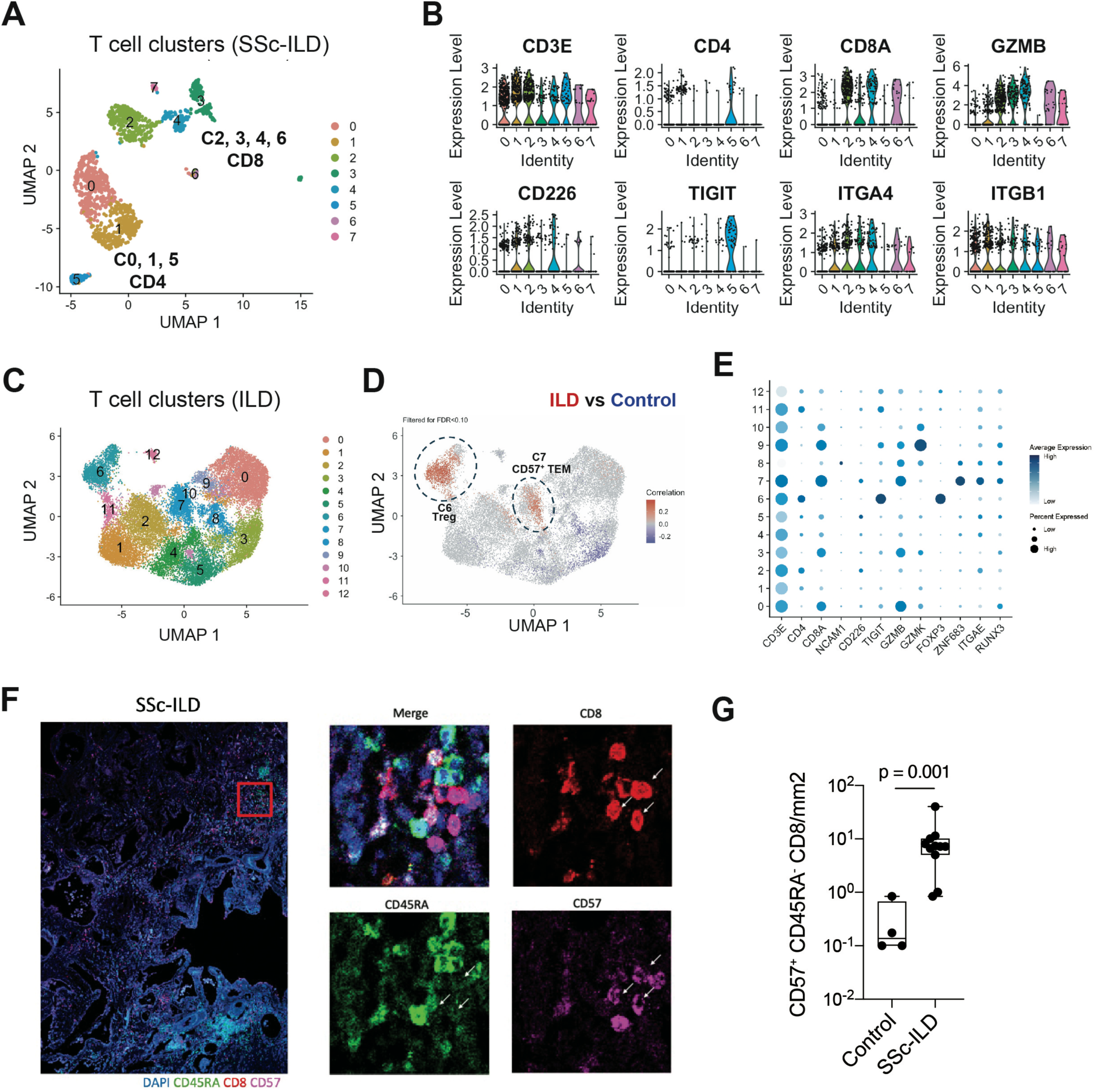
CD57^+^ TEM cells are expanded in SSc-ILD lung tissues. **A.** UMAP clustering of T cells in lung tissue scRNA-seq from 4 SSc-ILD donors (*23*). **B.** Gene expression of T cell clusters from (A). **C.** UMAP clustering of T cells in lung tissue scRNA-seq generated from 66 ILD and 48 control donors (*24*). **D.** CNA analysis of CD8 T cells from the scRNA-seq, adjusting for age and sex. Red indicates cell neighborhoods enriched in ILD samples. **E.** Gene expression of the CD8 clusters in the scRNA-seq data in (C). **F.** Representative images of CD57, CD8, CD45RA staining of SSc-ILD lung tissue. **G.** Quantification of CD57^+^ CD45RA^-^ CD8 in tissue staining. (HC: n=4, SSc-ILD: n = 11). Mann-Whitney U test.

### CD57^+^ TEM infiltrated and expanded in the patients with SSc-ILD

We next sought to determine whether CD57^+^ TEM cells are present within lungs of patients with SSc-ILD. We first analyzed published scRNA-seq data from lungs of 4 SSc-ILD patients (*23*), which demonstrated the presence a CD8 T cells expressing *GZMB* and *CD226* (Figure 4A, B), consistent with the phenotype expanded in circulation of SSc-ILD patients. To assess accumulation of CD8 T cells across a larger range of ILD samples, we analyzed a published lung tissue scRNA-seq dataset generated from 114 donors (66 ILD and 48 controls) (*24*). CNA analysis of T cell clusters indicated that C6 (Treg) and C7 (cytotoxic CD8 T cells resembling CD57^+^ TEM) were highly enriched in ILD lung samples compared to controls (Figure 4C, D). Interestingly, unlike CD57^+^ TEM in blood, C7 (CD57^+^ TEM) exhibited high expression of *ITGAE* (CD103) and *RUNX3* (Figure 5E), markers of Trm cells (*25*), suggesting that CD57^+^ TEM acquire Trm features in lung, allowing them to maintain residence in the tissue. To confirm the presence of CD57^+^ CD8 T cells in SSc-ILD lungs unambiguously at the protein level, we quantified CD57^+^ TEM by immunofluorescence microscopy in control donors, SSc-ILD, and idiopathic pulmonary fibrosis (IPF) samples collected via VATS or as lung explants. CD57^+^ TEM cells were abundant in lung tissue from SSc-ILD patients compared to controls (Figure 4F, G). Together, these results suggest that the CD8 T cells infiltrating SSc-ILD lung tissues share similar characteristics with the expanded population of CD57^+^ TEM in the circulation.

**Figure 5.**
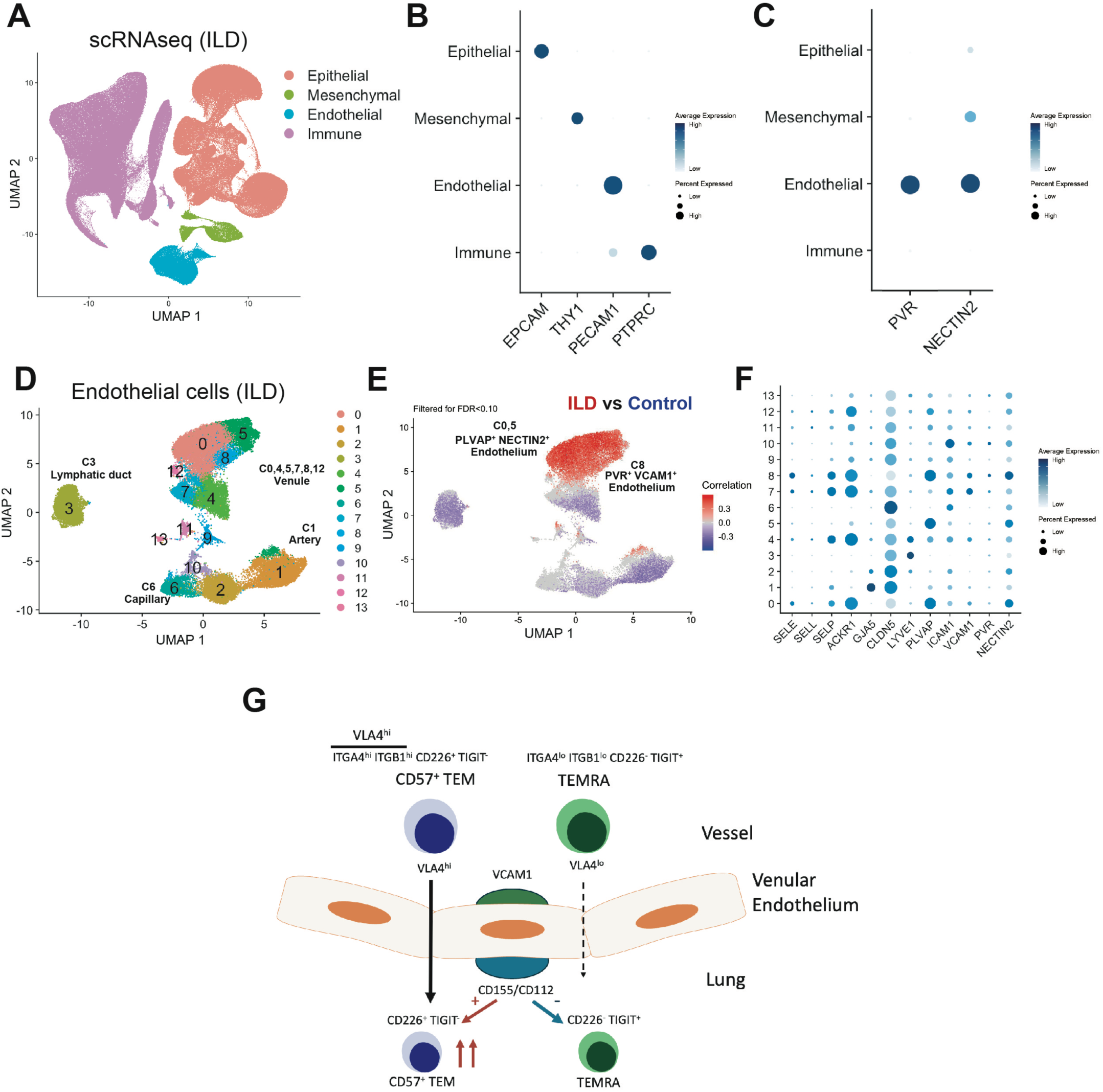
Endothelial cells are the main source of CD155 and CD112 in ILD tissues. **A.** UMAP clustering of all cells in lung tissue scRNA-seq generated from 66 ILD and 48 control donors (*24*). **B.** Expression of EPCAM, THY1, PECAM1, and PTPRC in cell subsets from (B).**C.** Expression of CD155/PVR and CD112/NECTIN2 in cell subsets from (B). **D.** UMAP clustering of endothelial cells in scRNA-seq data from (B). **E.** CNA analysis of endothelial cells from (B) comparing ILD vs control samples. Red indicates cell neighborhoods enriched in ILD samples. **F.** Expression of CD155/PVR and CD112/NECTIN2 in endothelial clusters. **G.** Schematic diagram of the infiltration and activation of CD57^+^ TEM in SSc-ILD pathogenesis.

### Expansion of CD155^+^ CD112^+^ VCAM1^+^ endothelial cells in an ILD cohort

Given evidence that ligands for CD226 augment activation of CD57^+^ TEM cells, we interrogated the expression of the ligands CD155 and CD112 in SSc-ILD lung tissue (Figure 5A). CD155 was detected in macrophages and vascular endothelial cells in lungs of patients with SSc-ILD (Supplementary Figure 6) (*23*). CD112/NECTIN2 was not detected in this dataset. In the larger dataset of cells across different ILD lung samples, both CD155/PVR and CD112/NECTIN2 were expressed highly in endothelial cells (Figure 5B, C) (*24*). CNA analysis of the broader ILD dataset focused on the endothelial cells revealed that three venule clusters, C0, C5, and C8, were highly enriched in ILD samples (Figure 5D, E). Notably, these three clusters highly expressed NECTIN2 and PLVAP. PLVAP forms a diaphragm that regulates vascular permeability (*26–28*). Interestingly, C8 also demonstrated high expression of CD155/PVR and VCAM1 (Figure 5F), the latter of which is a receptor for the integrin VLA4. A similar endothelial cell cluster highly expressing VCAM1 and CD155/PVR was also detected in IPF Cell Atlas (Supplementary Figure 7) (*29*, *30*). Given that CD57^+^ TEM cells highly express VLA4, the interaction between VLA4 on CD57^+^ TEM and VCAM1 on venular endothelial cells may promote the migration of CD57^+^ TEM into lung tissue. This migration, followed by CD155-mediated activation, may enhance the cytotoxic activity of CD57^+^ TEM (Figure 5G).

## Discussion

In this study, we used extensive mass cytometry and scRNA-seq profiling data from a well characterized cohort of patients with SSc to identify a unique cytotoxic CD8 T cell subset that is expanded in the circulation of SSc-ILD subjects and accumulates within SSc-ILD lungs. The expanded CD57^+^ TEM population is distinguished from TEMRA CD8 T cells by a distinct transcriptomic signature and high expression of VLA4 integrin and CD226, which may allow activation by and targeting of endothelial cells. Recognition of this expanded CD57^+^ TEM population with pathologic function provides a specific feature of immune activation in SSc-ILD that may serve as a therapeutic target and as a biomarker for SSc-ILD.

Here, we demonstrate that CD57^+^ CD8 TEM cells are expanded in multiple cohorts of patients with SSc-ILD. These findings are extend prior observations that CD57^hi^ CD4 CTLs are increased in patients with SSc, and among these patients, those with a higher proportion of CD57^hi^ CD4 CTLs have a higher prevalence of lung disease complications (*12*). Endothelial cells may be a potential key driver of the expansion of this immune cell subset. Our observation of high expression of VLA4 and CD226 on CD57^+^ TEM cells provides new evidence for a likely interaction between cytotoxic T cells and venular endothelial cells, which express both VCAM1 and CD155, as a target of the cytotoxic T cell response. Considering that blood flow velocity is high in arterial regions, where it is difficult for immune cells to adhere to the endothelium, it is reasonable that endothelial cells in the post-capillary area support the infiltration of CD57^+^ TEM cells. Escalante et al. reported that CD155 expression on HUVECs was enhanced by IFN-γ and IL-1 (*31*). This mechanism may act as an immune checkpoint for T cells expressing TIGIT, such as TEMRA cells. In contrast, CD57^+^ TEM cells with little TIGIT but high CD226 expression may evade CD155-dependent immunotolerance and instead exert pro-inflammatory effects via CD226 signalling. Given that the pathogenesis of ILD appears to be primarily driven by vascular endothelial damage, subsequent alveolar epithelial cell injury, maladaptive repair, and fibrosis, this activating effect of endothelial cells may enhance the initial process of cytotoxic injury to the endothelium mediated by CD57^+^ TEM cells.

One of the distinguishing features of CD57^+^ TEM is the high expression of ZNF683. ZNF683, also known as Hobit, is a key transcription factor for Trm cells, which reside in tissues and respond rapidly to viral and bacterial infections in the lungs (*32–34*). Interestingly, CD57^+^ TEM in lung tissue showed high expression of ITGAE (CD103) and RUNX3, which are characteristic of Trm cells. In contrast, CD57^+^ TEM in peripheral blood of patients with SSc-ILD did not express these markers. We hypothesize that circulating CD57^+^ TEM may acquire Trm-like characteristics upon infiltrating the lungs, allowing them to reside and manifest their effector cytotoxic function within affected tissues, thus contributing to pathogenetic injury and chronic inflammation in ILD. Furthermore, ZNF683-expressing CD8 T cells in blood have been reported as enriched for antigen reactivity in a case of kidney allograft rejection following immune checkpoint inhibitor therapy (*35*). The TCRs from alloreactive CD8 T cells infiltrating the rejected kidneys showed a high overlap with the TCRs of ZNF683-expressing CD8 T cells in blood. These findings further support the concept that circulating ZNF683-expressing CD8 T cells migrate to sites of inflammation, where they contribute to tissue damage.

The detection of expanded CD57^+^ TEM cells in the circulation of patients with SSc-ILD nominates this cellular feature as a potential biomarker to identify patients at risk for progressive ILD or to track the extent of the ongoing pathologic immune activation. The current study is limited by the lack of a prospective analysis of the association of CD57^+^ TEM cells with lung disease in SSc patients and the absence of paired blood and lung samples. Further, we do not know whether the expanded CD57^+^ TEM cells recognize relevant antigens in SSc and have not tested their function in an in vivo context. Nonetheless, the study highlights a specific, clonally expanded CD8 T cell population with plausible pathologic function that is strongly associated with SSc-ILD. As presence of ILD carries an important prognostic value not only in SSc but also in other autoimmune rheumatic conditions such as RA and myositis, it will be of great interest to determine whether CD57^+^ TEM cells are similarly increased in ILD patients affected by these conditions and whether this cellular subset can be targeted for therapeutic benefit.

## Acknowledgements

This work was supported in part by Department of Defense Scleroderma Research Program Translational Research Partnership Award SL200016P1 (to FB and DAR), Doris Duke Charitable Foundation Clinical Scientist Development Award, Burroughs Wellcome Fund Career Award in Medical Sciences, and NIAMS, NIH P30-AR-070253 award (to DAR), and the Kao Multidisciplinary Scleroderma Program at Cedars-Sinai (to FB).

## Author contributions

T.S. conceived the project, performed experiments, analyzed data, and wrote the manuscript. Y.C., R.A., and K.T. analyzed transcriptomic data. K.E.M. and K.H. assisted with the cytotoxicity assay. M.E. collected clinical information. N.B. participated in study design and data interpretation. P.W., M.K., and E.Y.K. collected tissue samples for histology. F.B. and D.A.R. conceived the project, supervised the work, recruited the patients, and wrote the manuscript. All authors discussed the results and revised the manuscript.

## Method

### Sample collection

Peripheral blood mononuclear cells (PBMC) were collected from 67 patients with SSc at the University of California, San Francisco (UCSF), and from 15 patients with SSc and 18 healthy controls from Brigham and Women’s Hospital (BWH). All SSc patients fulfilled the 2013 American College of Rheumatology/EULAR criteria for SSc (*36*). The study was approved by the IRB at UCSF and BWH (UCSF IRB# 15-16463, BWH Protocol #: 2014P002558). All patients were consented to collect blood samples and clinical information. The presence of ILD was confirmed by radiographic evidence of pulmonary fibrosis on high-resolution computed tomography (HRCT) conducted in patients exhibiting forced vital capacity (FVC) <80%, absolute decline of the FVC (L) >10% over two consecutive assessments, or with evidence of bibasilar crackles at lung auscultation. Pulmonary arterial hypertension (PAH) was confirmed with right heart catheterization by mean pulmonary artery pressure (mPAP) >25 mm Hg, peripheral vascular resistance >3 Wood units, and pulmonary capillary wedge pressure <15 mmHg as the updated 2022 haemodynamic definition of PAH was not yet established at the time of patient enrolment (*37*). Healthy controls were screened for any evidence of autoimmune systemic disease, neoplasm, or lung pathology. Blood samples were collected into heparin tubes, and PBMC was isolated by density centrifugation using Ficoll-Hypaque in 50-mL conical tubes. PBMC was cryopreserved in 10% DMSO-containing solution.

### Mass cytometry

Cryopreserved PBMC was thawed into complete culture media (RPMI 1640 Medium supplemented with 5% heat-inactivated fetal bovine serum, 1 mM GlutaMAX, 10 mM HEPES, and penicillin-streptomycin). Cells were counted, and 0.2 × 10^6^ to 2 × 10^6^ cells from each sample were transferred to a 96-well conical bottom polypropylene plate for staining. After the cells were transferred to the plate, the viability staining reagent cisplatin with a dilution of 1:1,000 was added to the cells directly for two minutes and then diluted with culture media. After centrifugation, a human Fc receptor blocking agent was added at a 1:100 dilution with cell staining buffer for 10 minutes, followed by incubation with metal-conjugated antibodies for 30 minutes at room temperature. Antibodies were obtained from the Harvard Medical Area cytometry by time-of-flight mass spectrometry (CyTOF) Antibody Resource and Core and Fluidigm. Cells were then fixed in 4% paraformaldehyde for 10 minutes before permeabilization with the FoxP3/Transcription Factor Staining Buffer Kit (eBioscience). Cells were incubated in Transcription Factor Fix/Perm Buffer for 30 minutes before barcoding. Cells were then barcoded using the Cell-ID 20-Plex Pd Barcoding kit (Fluidigm) and pooled together into one tube. The metal-conjugated intracellular antibody mix was then added into the tube, and cells were incubated for 30 minutes. Cells were then fixed with 4% paraformaldehyde for 10 minutes and then washed out and resuspended in a cell staining buffer and left at 4 degrees overnight. The next day, DNA was labeled for 20 minutes with iridium intercalator solution (Fluidigm). Samples were washed and then counted in the presence of EQ Four Element Calibration beads at a final concentration of 1 × 10^6^/mL. Samples were acquired on a Helios CyTOF Mass Cytometer. Mass cytometry data were generated on PBMC samples from patients with SSc and healthy controls using a 39-marker mass cytometry panel designed to identify T cell subsets. Samples were processed in five batches, with 20 barcoded samples in each batch. Patient samples were analyzed in barcoded batches, with each batch randomized to include samples from both healthy controls and patients with SSc with different degrees of lung involvement. The raw flow cytometry standard files were normalized together using bead standard normalization to minimize batch effects and were debarcoded for analysis.

### Single-cell RNA sequencing

The files were processed with the Seurat R package (v5.1.0) (*38*). After QC, mRNA expression was log-normalized (scale.factor = 10,000). Then the highly variable genes, selected through variance stabilizing transformation, were scaled. Next, principal components (PCs) were calculated based on mRNA data, and samples were batch-corrected using the Harmony R package. UMAP visualization and clustering were performed. Sequencing reads for the TCR library were processed through the Cell Ranger workflow. Reads were aligned to a TCR reference (vdj-GRCh38), productive contigs were filtered, and CDR3 sequences were identified for each cell. Cells with matching both CDR3alpha and CDR3beta were grouped into the same clonotype. Each cell’s TCR was then paired with its transcriptomic data by matching cell barcodes. These files were then analyzed using the scRepertoire package (*39*).

### Flow cytometry

After cryopreserved PBMC was thawed into complete culture media, PBMC was washed with PBS and was stained with violet viability dye (ThermoFisher) on ice for 15min. After washing with PBS, cells were stained with flow antibodies on ice for 30min. When performing intracellular staining, cells were fixed with Foxp3 Fixation/Permeabilization (eBioscience) for 30 min at room temperature. After washing with Permeabilization Buffer (eBioscience), cells were stained with anti-GZMB antibody and anti-GZMK antibody on ice for 30min. After washing twice in 1x eBioscience Permeabilization Buffer, and passing through a 70μM filter, data were acquired on a BD Fortessa analyzer using FACSDiva software and analyzed using FlowJo.

### Bulk RNA sequencing

After cryopreserved PBMC from 5 SSc-ILD donors was thawed into complete culture media, PBMC was washed with PBS and was stained with violet viability dye (ThermoFisher) on ice for 15min. After washing with PBS, cells were stained with flow antibodies on ice for 30min. After washing with 1% BSA in PBS, approximately 20,000 cells of naive CD8 (Live^+^ CD3^+^ CD8^+^ CD56^-^ CD57^-^ CD27^+^ CCR7^+^ CD45RA^+^), CD57-TEM (Live^+^ CD3^+^ CD8^+^ CD56^-^ CD57^-^ CD27^+^ CCR7^-^ CD45RA^-^), CD57^+^ TEM (Live^+^ CD3^+^ CD8^+^ CD56^-^ CD57^+^ CD27^-^ CCR7^-^ CD45RA^-^), and TEMRA CD8 T cells (Live^+^ CD3^+^ CD8^+^ CD56^-^ CD57^+^ CD27^-^ CCR7^-^ CD45RA^+^) T cell populations were directly sorted using a 5L BD FACSAria Fusion cell sorter into RLT lysis buffer kept on ice. Cells were sorted through an 85μM nozzle. After extracting RNA using RNeasy Kit (Qiagen #74104), the samples were submitted to Molecular Biology Core Facilities at Dana-Farber Cancer Institute, and then generated data using mRNAseq SS v4. Raw reads were mapped using the STAR method protocol (https://github.com/alexdobin/STAR). Raw read counts were analyzed by R (Ver. 4.2.0) using DESeq2. After normalising RNA expression PCA was performed using prcomp and plotted with ggtools2. Differential gene expression was carried out using DESeq2 and volcano plots generated using ggplot2.

### Tissue CD57^+^ TEM staining

Unstained FFPE-fixed lung tissue samples were obtained from the BWH pathology core and USCF. After deparaffinization with D-limonene (3min twice) and rehydration with 100% (3min), 95% (3min), and 70% (3min twice) ethanol, the antigen retrieval was performed using Antigen Retrieval Buffer (ab93678) at 95 degrees for 10 min. After blocking with PBS, 3% BSA for 30min, the cells were stained with CD57 (x100), CD8 (x25), CD45RA (x100) primary antibodies in 3% BSA + 1% Triton X for 1 hour at room temperature. After washing with PBS, the slides were stained with AF488 (x500), AF555 x250, and AF647 x500 for 1 hour at room temperature. After washing with PBS, DAPI Fluoromount-G (Southern Biotech 0100-20) was mounted on the slides, and the samples were covered by the cover glasses. The immunofluorescence images were taken by Zeiss LSM 800, and then analyzed by QuPath (*40*).

### Cytotoxicity Assays

CD8 T cells were isolated from collars with MACS total CD8 isolation kit (Miltenyi) and subsequently sorted with a 5-laser BD FACSAria Fusion cell sorter for CD57^+^ TEM and TEMRA. CD32-expressing L cells were incubated with an agonist anti-CD3 antibody (SK7, Thermofisher) for 30 min on ice and then plated with sorted CD8 T cells at a ratio of 3:1 CD8 T cells:L cells. After 10 hours, the T cells and L cells were stained with AnnexinV APC and 7-AAD in Annexin Binding Buffer (all from Biolegend) and analyzed on a BD CantoII analyzer.

---

**Supplementary Figure 1.**
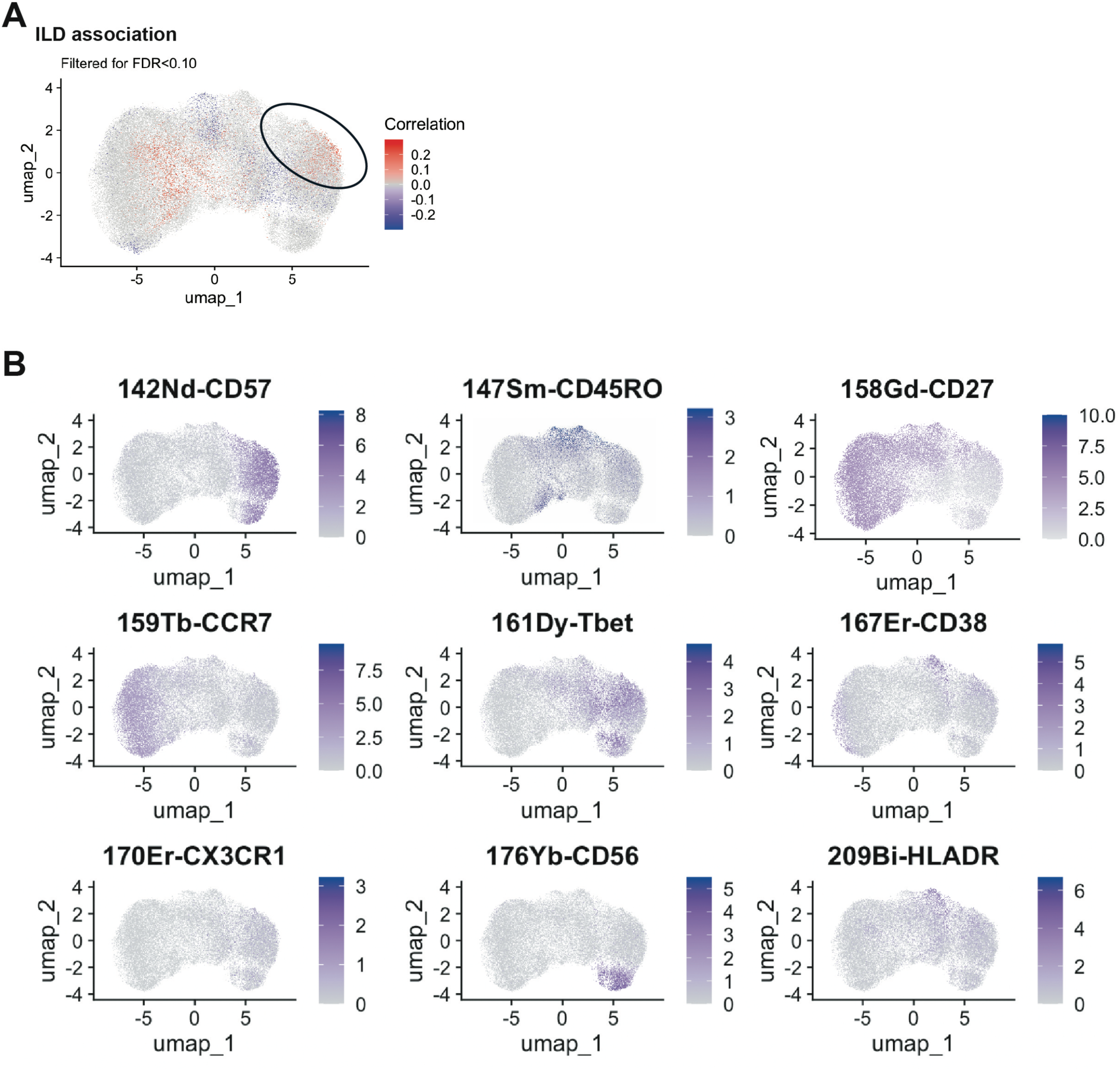
Marker expression of CD57^+^ TEM in mass cytometry. **A.** CNA analysis using CD8 T cells from HC and SSc-ILD patients, adjusting for age and sex. Red indicates cell neighborhoods enriched in SSc-ILD patients. **B.** Marker expression in T cells in mass cytometry data.

**Supplementary Figure 2.**
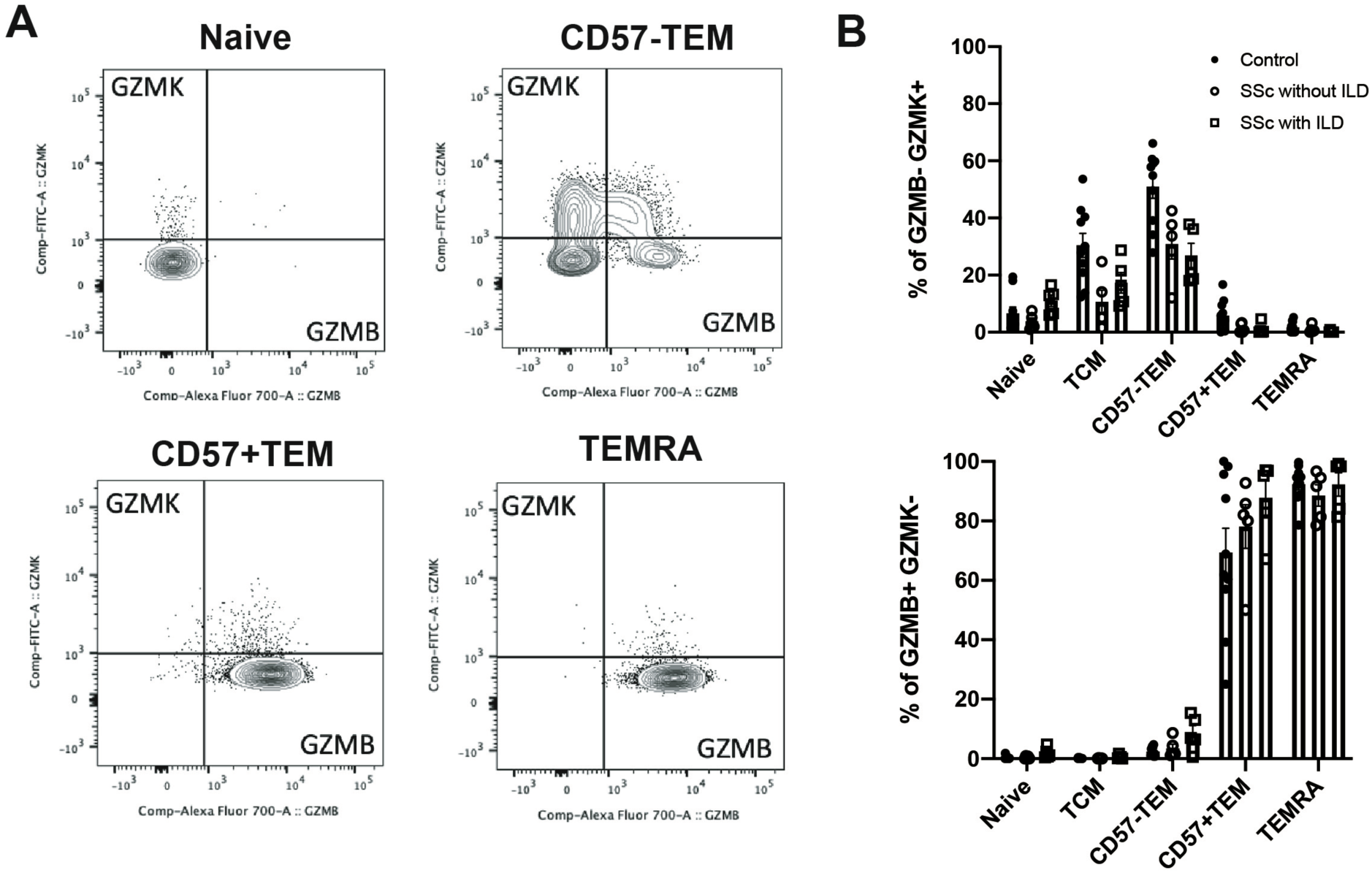
CD57^+^ TEM and TEMRA CD8 T cells are granzyme B producers. **A, B.** Intracellular staining of granzyme B and granzyme K in naive, TCM, CD57^-^ TEM, CD57^+^ TEM, and TEMRA CD8 T cells. HC: n = 10, SSc-non ILD: n = 5, SSc-ILD: n = 5.

**Supplementary Figure 3.**
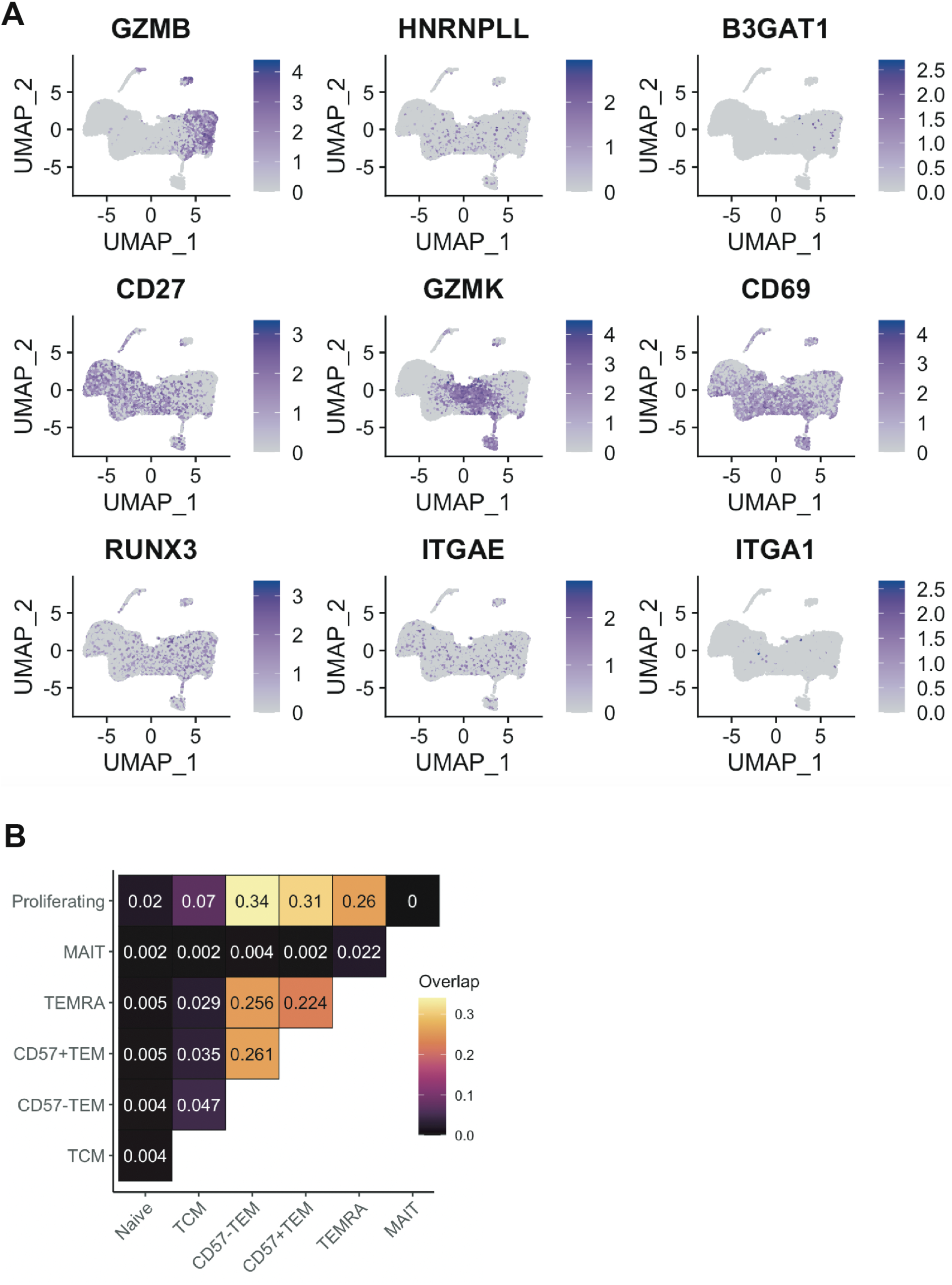
Gene expression of CD57^+^ TEM in scRNAseq data. **A.** Gene expression of *GZMB*, *HNRNPLL*, *B3GAT1*, *CD27*, *GZMK*, *CD69*, *RUNX3*, *ITGAE*, and *ITGA1* in scRNA-seq dataset from PBMC. **B.** TCR clonal overlap across indicated CD8 T cell clusters.

**Supplementary Figure 4.**
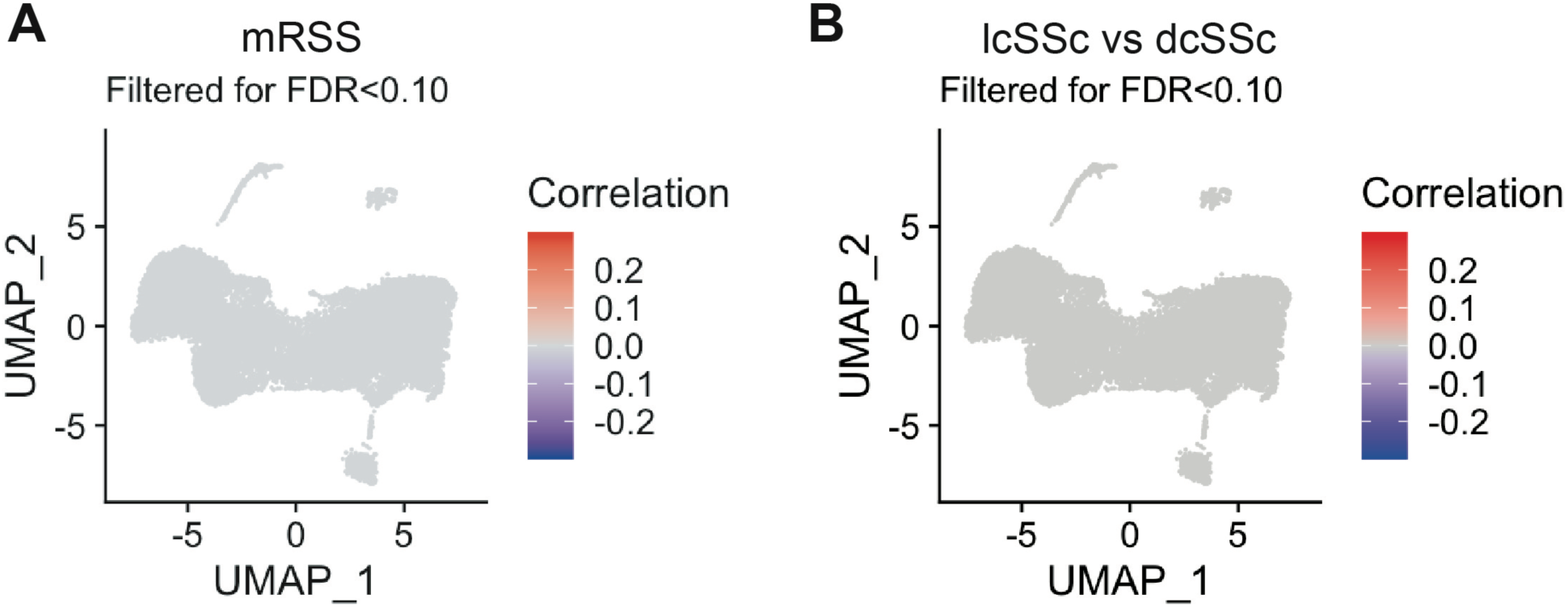
There is no association between CD57^+^ TEM and skin involvement. **A, B.** CNA analysis to evaluate the association between CD8 T cell cluster and severity of skin involvement, adjusting for age and sex.

**Supplementary Figure 5.**
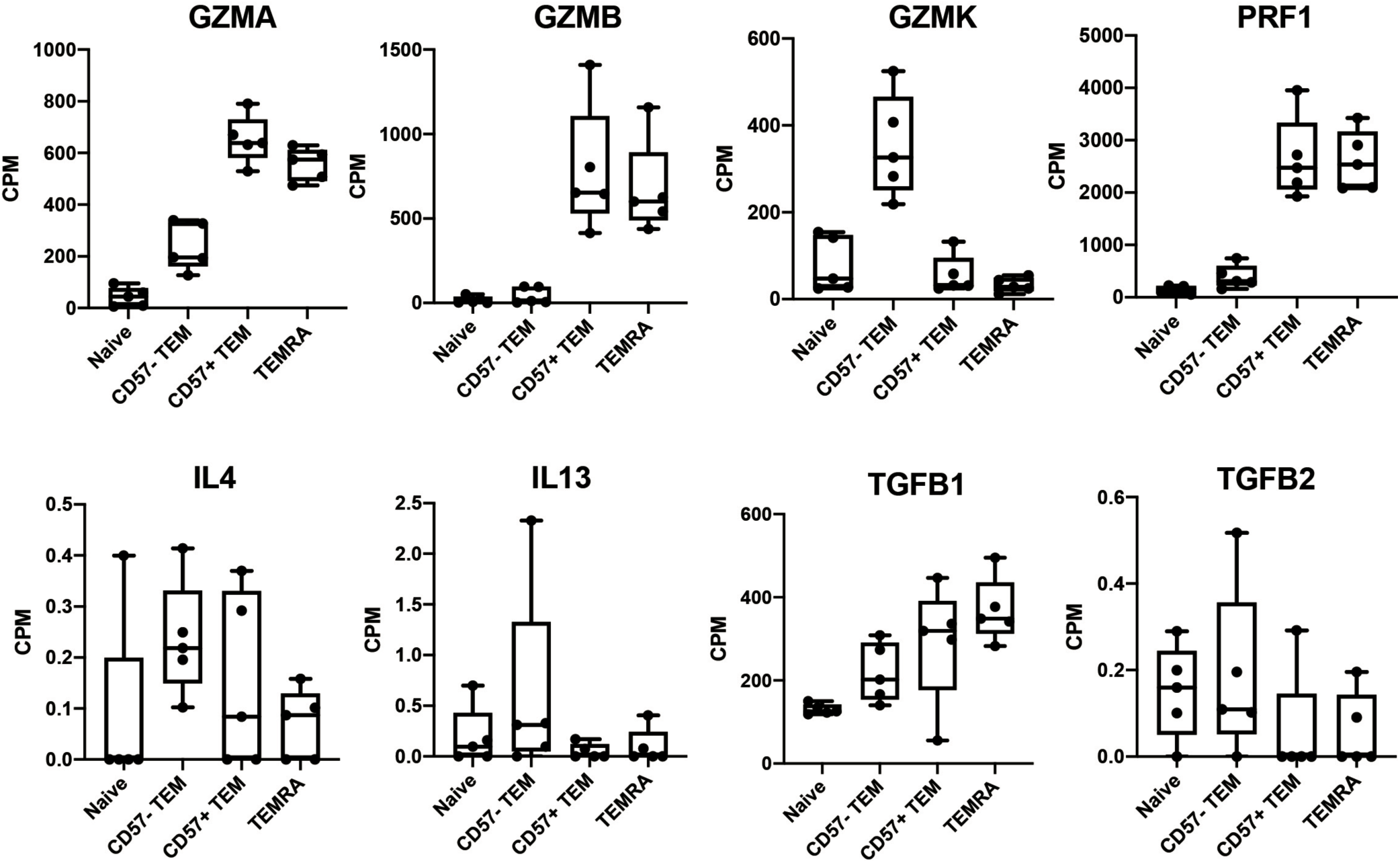
Gene expression in the bulk RNA-seq. Gene expression of *GZMA*, *GZMB*, *GZMK*, *PRF1*, *IL4*, *IL13*, *TGFB1*, and *TGFB2* in naive, CD57^-^ TEM, CD57^+^ TEM, and TEMRA CD8 T cells sorted from 5 SSc-ILD donors. CPM: Counts per million.

**Supplementary Figure 6.**
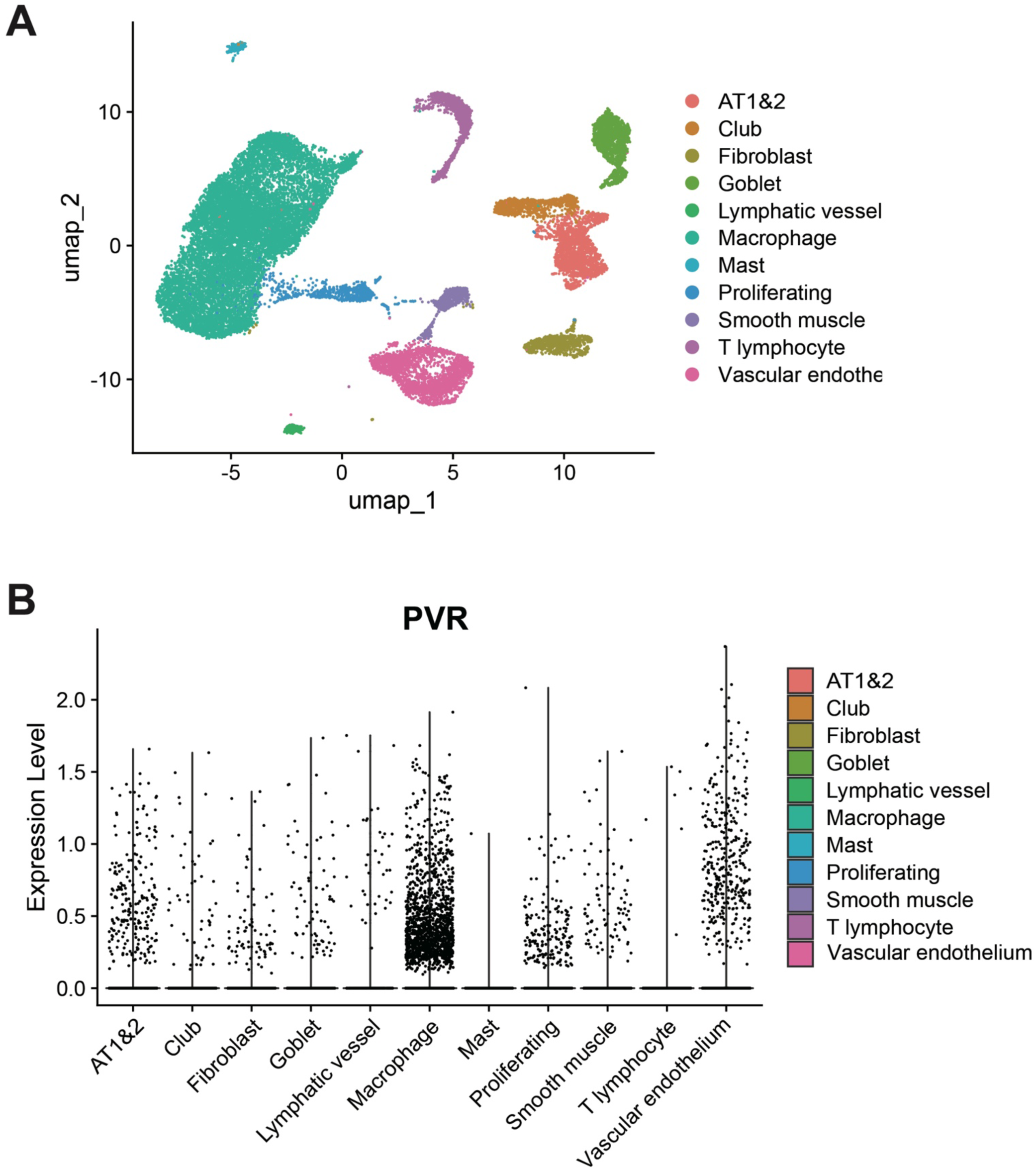
CD155/PVR expression in lung tissues from SSc-ILD. **A.** UMAP clustering of all cells in scRNA-seq from lung samples of 4 SSc-ILD donors (*23*). **B.** *CD155/PVR* expression in clusters from scRNA-seq data.

**Supplementary Figure 7.**
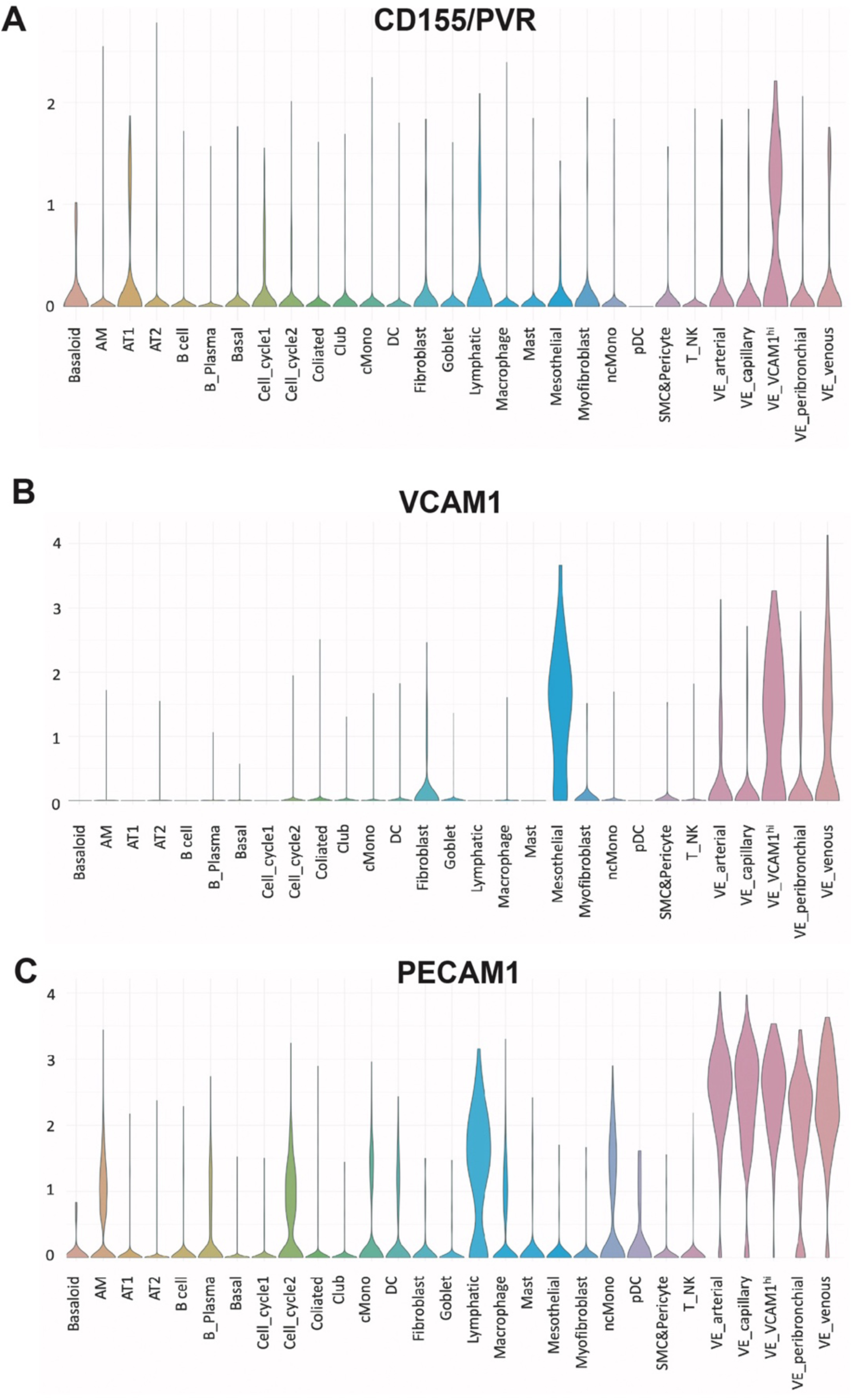
CD155/PVR expression in lung tissues from IPF Cell Atlas. **A-C.** *CD155/PVR*, *VCAM1*, and *PECAM1* expression in clusters from the IPF Cell Atlas scRNA-seq data (*29*, *30*).

**Supplementary Table 1.**
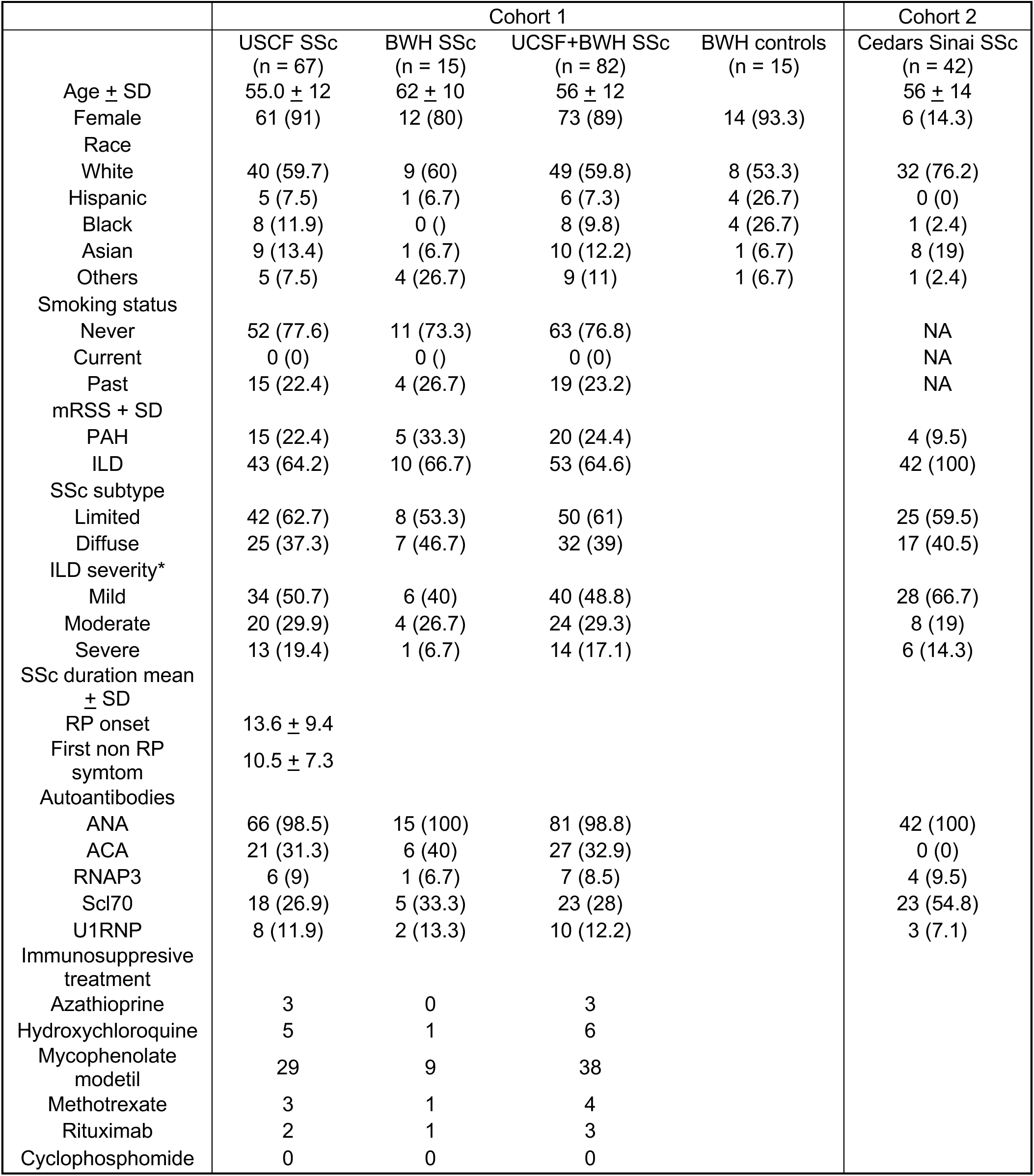
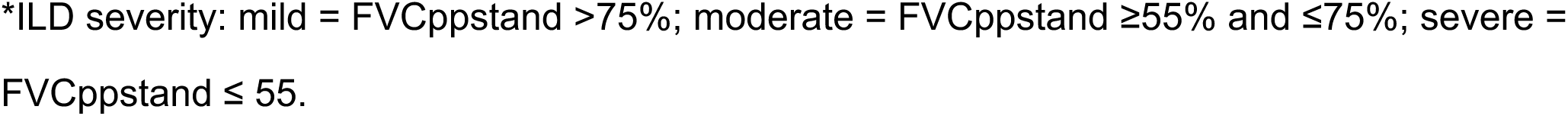
Demographics and clinical characteristics of patients in Cohort 1 and Cohort 2.

|  | Cohort 1 |  |  |  | Cohort 2 |
| --- | --- | --- | --- | --- | --- |
|  | USCF SSc<br>(n = 67) | BWH SSc<br>(n = 15) | USCF+BWH SSc<br>(n = 82) | BWH controls<br>(n = 15) | Cedars Sinai SSc<br>(n = 42) |
| Age ± SD | 55.0 ± 12 | 62 ± 10 | 56 ± 12 | | 56 ± 14 |
| Female | 61 (91) | 12 (80) | 73 (89) | 14 (93.3) | 6 (14.3) |
| Race |  |  |  |  |  |
| White | 40 (59.7) | 9 (60) | 49 (59.8) | 8 (53.3) | 32 (76.2) |
| Hispanic | 5 (7.5) | 1 (6.7) | 6 (7.3) | 4 (26.7) | 0 (0) |
| Black | 8 (11.9) | 0 ( ) | 8 (9.8) | 4 (26.7) | 1 (2.4) |
| Asian | 9 (13.4) | 1 (6.7) | 10 (12.2) | 1 (6.7) | 8 (19) |
| Others | 5 (7.5) | 4 (26.7) | 9 (11) | 1 (6.7) | 1 (2.4) |
| Smoking status |  |  |  |  |  |
| Never | 52 (77.6) | 11 (73.3) | 63 (76.8) |  | NA |
| Current | 0 (0) | 0 ( ) | 0 (0) |  | NA |
| Past | 15 (22.4) | 4 (26.7) | 19 (23.2) |  | NA |
| mRSS ± SD | | | | | |
| PAH | 15 (22.4) | 5 (33.3) | 20 (24.4) |  | 4 (9.5) |
| ILD | 43 (64.2) | 10 (66.7) | 53 (64.6) |  | 42 (100) |
| SSc subtype |  |  |  |  |  |
| Limited | 42 (62.7) | 8 (53.3) | 50 (61) |  | 25 (59.5) |
| Diffuse | 25 (37.3) | 7 (46.7) | 32 (39) |  | 17 (40.5) |
| ILD severity* |  |  |  |  |  |
| Mild | 34 (50.7) | 6 (40) | 40 (48.8) |  | 28 (66.7) |
| Moderate | 20 (29.9) | 4 (26.7) | 24 (29.3) |  | 8 (19) |
| Severe | 13 (19.4) | 1 (6.7) | 14 (17.1) |  | 6 (14.3) |
| SSc duration mean<br>± SD | | | | | |
| RP onset | 13.6 ± 9.4 | | | | |
| First non RP<br>symptom | 10.5 ± 7.3 | | | | |
| Autoantibodies |  |  |  |  |  |
| ANA | 66 (98.5) | 15 (100) | 81 (98.8) |  | 42 (100) |
| ACA | 21 (31.3) | 6 (40) | 27 (32.9) |  | 0 (0) |
| RNAP3 | 6 (9) | 1 (6.7) | 7 (8.5) |  | 4 (9.5) |
| Scl70 | 18 (26.9) | 5 (33.3) | 23 (28) |  | 23 (54.8) |
| U1RNP | 8 (11.9) | 2 (13.3) | 10 (12.2) |  | 3 (7.1) |
| Immunosuppressive<br>treatment |  |  |  |  |  |
| Azathioprine | 3 | 0 | 3 |  |  |
| Hydroxychloroquine | 5 | 1 | 6 |  |  |
| Mycophenolate<br>modetil | 29 | 9 | 38 |  |  |
| Methotrexate | 3 | 1 | 4 |  |  |
| Rituximab | 2 | 1 | 3 |  |  |
| Cyclophosphomide | 0 | 0 | 0 |  |  |

\*ILD severity: mild = FVCppstand >75%; moderate = FVCppstand ≥55% and ≤75%; severe = FVCppstand ≤ 55.

